# Multiple protein-protein interactions drive the assembly and budding of the Chikungunya virion

**DOI:** 10.1101/2025.10.10.681735

**Authors:** Miguel Angel Coronado-Ipiña, Shuyang Zhang, Siyu Li, Pedro Gerardo Hernández-Sánchez, Denisse Cadena-Lopez, Vanessa Labrada-Martagón, Ajay Gopinathan, Suchetana Mukhopadya, Roya Zandi, Mauricio Comas-Garcia

## Abstract

The assembly of enveloped viruses is a highly orchestrated process that depends on the coupling of multiple protein–protein interactions within a membrane environment. To gain mechanistic insight into this process, we use Chikungunya virus as a model system to study Alphavirus assembly, focusing on the interplay between core–spike and spike–spike interactions. We begin with coarse-grained molecular dynamics simulations to systematically explore how the symmetry of the nucleocapsid core, together with the relative strengths of spike–core and spike– spike interactions, influences budding efficiency and the emergence of icosahedral particle symmetry. Building on these computational predictions, we perform site-directed mutagenesis on Chikungunya virus 181/25 and examine the consequences for particle assembly and budding in cultured cells, as well as the impact of these mutations during *in-cellulo* assembly. Our results reveal that canonical core–spike interactions, while necessary, are not sufficient for successful assembly. Instead, lateral interactions among glycoproteins emerge as critical determinants of efficient budding, particle stability, and the maintenance of icosahedral symmetry. Together, these findings provide an integrated computational and experimental framework for understanding the molecular principles governing Alphavirus assembly.

---

The assembly of enveloped viruses is a complex process orchestrated by protein–protein and protein–membrane interactions to drive membrane deformation, budding, and packaging of the viral genome into a fully infectious particle^1–7^. For certain viruses, this process is known to rely primarily on matrix or capsid proteins that recruit host-cell machinery to mediate budding from the plasma membrane ^6, 8–13^. In contrast, the mechanisms of assembly and budding in Alphaviruses remain less well characterized, with some conflicting models ^2, 14–18^.

Alphaviruses are arthropod-borne viruses infecting diverse animal species, classified as New- or Old-World depending on whether they are endemic to the Americas or other regions ^19^. Chikungunya virus (CHIKV), an Old-World Alphavirus, is the most prevalent Alphavirus infecting humans and has caused recurrent epidemics of febrile illness and chronic polyarthritis worldwide^20–25^. Alphaviruses have a spherical virion approximately 70 nm in diameter, consisting of two concentric icosahedral protein shells separated by a lipid bilayer derived from the host plasma membrane. The outer shell comprises 240 copies of E1–E2 heterodimers arranged as 80 trimeric spikes, while the inner shell, nucleocapsid or core (NC), consists of 240 capsid proteins (CP) encapsidating the ∼12 kb single-stranded positive-sense RNA genome^14^. These two layers are structurally coupled via interactions between the cytoplasmic tail of E2 and a hydrophobic pocket in the capsid protein^14, 26^.

Cryo-electron microscopy (cryo-EM), cryo-electron tomography (cryo-ET), biochemical analysis, and molecular dynamics (MD) simulation studies of Alphaviruses have revealed four major classes of interactions stabilizing the virion: (1) intra-dimer interactions^3, 4^, (2) inter-dimer E1–E2^2, 16, 27–29^, (3) inter-spike trimer–trimer interactions^30, 31^, and (4) vertical contacts between E2 and the capsid core^26, 30, 32–34^. These interactions coordinate virion formation—from initiating budding and guiding icosahedral assembly to controlling particle yield, stability, and infectivity^14, 35, 36^ —though their precise roles and mechanisms remain unclear.

A central question in Alphavirus biology is how these two icosahedral layers assemble coordinately during budding. Several models have been proposed: in the assembly-from-within model, a preformed icosahedral core serves as a structural template for glycoprotein assembly ^15, 17^; in the assembly-from-without model, glycoproteins initiate budding and recruit the capsid core *post hoc*^16^; and in the concerted assembly model, an initially irregular cytoplasmic core gains order upon interacting with glycoproteins at the membrane^2, 18^. The latter has gained support from recent cryo-ET studies, which show that cores lack icosahedral symmetry in the cytoplasm but acquire structural order during budding ^2, 18^. These studies did not observe a pre-assembled lattice of spikes at the membrane or icosahedral cytoplasmic cores, suggesting that E2–core binding and lateral glycoprotein interactions drive budding and shape the geometry of the final virion. However, in the absence of direct, molecular-resolution visualization of both the budding process and spike organization under budding-inhibited conditions in infected cells, the mechanisms underlying icosahedral assembly and its inhibition remain poorly defined.

To understand the factors contributing to Sindbis (Alphavirus) budding, Hagan and co-workers developed a coarse-grained simulation model that incorporates spike–spike and spike– core interactions^30^. They showed that while glycoprotein interactions alone can drive budding, the resulting particles tend to be polydisperse and irregular. Even modest interactions with a rigid core improve uniformity and reduce kinetic trapping. However, their focus was on the conditions that permit or impede budding, rather than on achieving icosahedral symmetry in the final product.

Here, we identify the molecular determinants that govern CHIKV budding and assembly by pinpointing the specific interactions that coordinate these processes. One question that naturally arises is whether capsid proteins (CPs) preassemble into icosahedral nucleocapsids (NCs) that act as structural templates, or whether glycoprotein spikes instead drive the coordinated formation of both the outer spike lattice and the inner NC shell. To address this, we performed coarse-grained MD simulations to examine how spike–core and spike–spike interactions, together with core symmetry, influence budding efficiency and the emergence of icosahedral order. In particular, we tested whether immature, non-icosahedral NCs can serve as imperfect scaffolds that promote the formation of icosahedral spike shells.

In our MD simulations, we find that when the core lacks icosahedral symmetry and is constrained to a fixed geometry that prevents local dissociation and reassociation, the spikes fail to assemble into a complete icosahedral shell. Nevertheless, the absence of an entire pentamer does not completely abolish symmetry; spike assembly can still progress toward icosahedral order, albeit with reduced efficiency. These results strongly support a concerted assembly mechanism, in which tightly coordinated interactions across the membrane synchronize the formation of both the inner nucleocapsid and the outer spike lattice.

Guided by these insights, we conducted targeted mutational experiments using a CHIKV reporter virus to probe the functional roles of key glycoprotein interfaces. Specifically, we introduced E2 Y400K/L402R to disrupt spike–core interactions, E2 N263Q to weaken intertrimer contacts, and E1 K245N/R247T to perturb intradimer interactions. Our combined computational and experimental findings demonstrate that E2–core interactions are essential for initiating membrane curvature and nucleating particle formation, while strong lateral spike–spike contacts are required for the completion and stabilization of fully formed virions. Thus, both vertical (spike–core) and lateral (spike–spike) interactions act cooperatively to establish and maintain icosahedral symmetry across the two structural layers of the virus. This cooperative mechanism ensures proper virion architecture and infectivity and highlights potential antiviral targets aimed at disrupting envelope-protein organization to block particle assembly or release.

## RESULTS

Motivated by several experiments on CHIKV budding and the formation of icosahedral symmetry, carried out by Chiu’s group ^2, 18, 37^ and others^27, 30, 38–40^, we performed molecular dynamics (MD) simulations to investigate how specific spike–spike and spike–core interactions govern virus assembly. This approach isolates the effects of individual interactions in a controlled, simplified environment, and the resulting insights guided targeted mutational experiments using CHIKV 181/25 ^41^ to test their roles in particle formation in cultured cells— results presented in the sections following the simulations.

### Simulations

We simulated the budding of a nucleocapsid through a membrane that, as described in the *Methods and Materials* section, exhibits fluid behavior^41^ and contains inserted spike proteins (Figure 1A). Each spike comprises three domains: the ectodomain (ED), transmembrane domain (TMD), and cytoplasmic domain (CTD). The nucleocapsid is represented by a core with different geometries, illustrated in Figures 1B–1D, which can interact with the membrane spikes.

**Figure 1.**
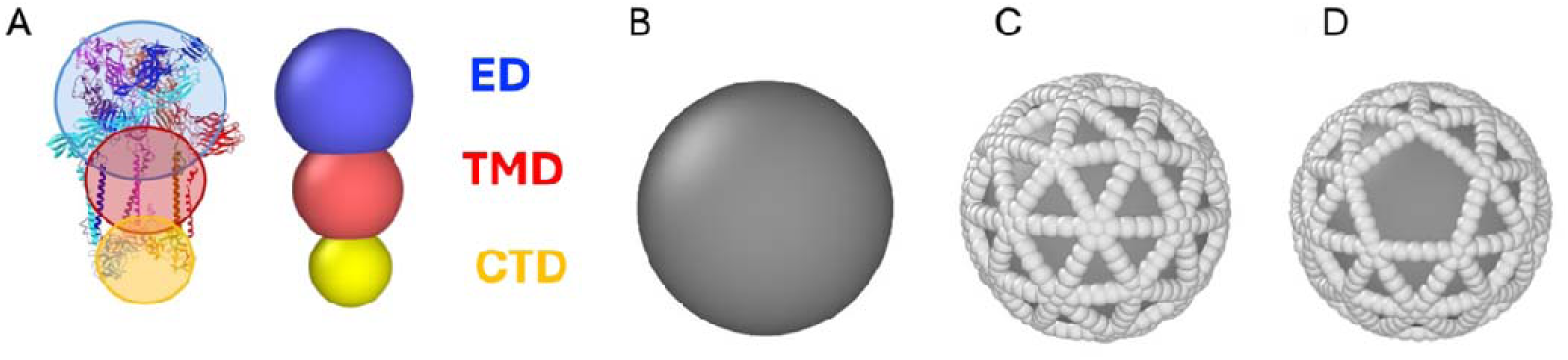
Schematic representation of a coarse-grained spike protein and core. A) The atomic structure of the CHIKV spike protein, a trimer of heterodimers (PDB ID: 8FCG). The shaded circles represent th components of the coarse-grained model, which consists of three spherical particles: ED (ecto domain, blue), TMD (transmembrane domain, red), and CTD (cytoplasmic domain, yellow). The size asymmetry between the domains allows the spike proteins to adopt a curved configuration, promoting membrane bending. B) A simple spherical core. C) A spherical core decorated with 80 triangular units arranged with icosahedral symmetry. D) A modified version of the core in which several triangles are missing, breaking the icosahedral symmetry.

Spike–spike interactions alone were first tested to determine whether they could drive budding in the absence of a core. Consistent with earlier work ^29, 30, 42, 43^, these interactions were sufficient to induce budding but produced irregular, non-icosahedral structures. Introducing a spherical core that interacts attractively with the spike cytoplasmic tails (see *Methods and Materials* for potential details) improved local ordering but failed to yield global icosahedral symmetry(see Supplementary Figure 3). Only when a preassembled icosahedral core was incorporated (Figure 1C) did the system produce a symmetric, virus-like shell.

In this configuration, three interaction classes determined assembly behavior: (i) CTD– core attraction (ε_mc_), (ii)TMD–TMD lateral association (ε_mm_), and (iii) excluded-volume repulsion among spike ectodomains (ED–ED). By systematically varying ε_mc_ and ε_mm_ we identified conditions that maximize budding efficiency and yield near-perfect icosahedral structures. The resulting trends were compared to experiments using CHIKV 181/25 infection and transfection of HEK-293T cells expressing a CMV-driven reporter gene. We note that mutations at the intradimer interface were introduced experimentally and shown to affect heterodimer interactions within a trimer; however, these were not modeled in our simulations, which treat the trimer as the smallest structural unit in a coarse-grained framework.

### Parameter Optimization for the Wild-Type System

Systematic exploration of the parameter space revealed an optimal regime corresponding to a CTD–core interaction strength of ε_mc_ = 74 k_B_T, together with the default ED size and TMD– TMD attraction (values listed in the figure caption). This condition, designated as the wild-type (WT) baseline, yields an effective E2–core interaction of approximately 8 k_B_T, consistent with previous studies^30^. Under these conditions, budding proceeds efficiently, with about 80±5 spike assemblies on the core to form a nearly icosahedral shell (Figure 2A). The inset in Figure 2B highlights the final spike distribution (yellow spheres), and Figure 2C presents a cutaway view showing their spatial arrangement around the core. Although the model lacks an explicit scission mechanism, budding terminates naturally once the shell closes around the core.

**Figure 2.**
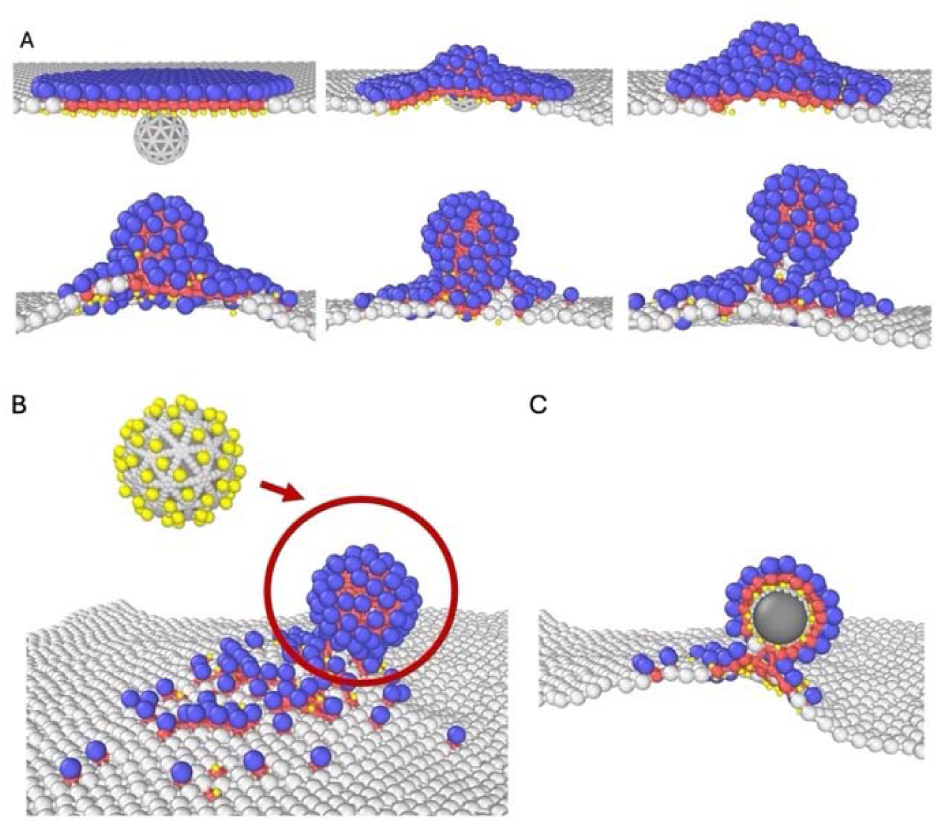
Snapshots of the budding process. A) Snapshots of the budding process using an icosahedral core at timesteps t=0,5,10, 80,250, and 700 (left to right, top to bottom). The system parameters include a CTD–core interaction strength of 74 kBT and a TMD-TMD interaction strength of 0.2 kBT. The particle diameters are set to 1.15 for ED, 0.8 for TMD, and 0.4 for CTD. B) Final state of the system after budding. The inset highlights the location of CTD particles along with the core. C) Cross-sectional view of the final state through the center of the assembled structure.

At weaker interaction strengths, however, this behavior changes markedly. When ε_mc_ falls below a critical threshold, the core fails to bud from the membrane—consistent with previous observations ^17, 26, 28, 32, 33^. In this regime, the membrane remains unbudded (see Supplementary Figure 4), reproducing the phenotype observed experimentally for the CT mutant described below.

### Modeling the Effects of Mutations by Tuning Attractive and Steric Interactions Between Spike Protein Domains

Ribeiro-Filho *et al*. proposed that in the Mayaro virus, the glycosylation site equivalent to CHIKV E2 N263 acts as a “*molecular handshake*” that stabilizes the viral particles^31^. This glycosylation could influence trimer–trimer interactions through steric repulsions or hindrances, and its removal may alter the local curvature or relative orientation between adjacent spikes, thereby making budding more difficult.

To quantify the absence of this glycosylation site in E2, we conducted a series of simulations with icosahedral cores and compared their relative success rates. To isolate the effects of specific mutations, we introduced changes by modifying one or more of these parameters, while holding the CTD–core interaction constant at ε_mc_ = 74 k_B_T, same as the WT system interaction strength. For each condition, we performed eight independent simulation trials. In the WT system (ε_mc_ = 74 k_B_T), 7 out of 8 simulations resulted in successful assemblies, which we define as a 100% baseline success rate. A configuration was considered successful if no more than five of the 80 spike proteins, expected in a perfect T = 4 structure, were missing or mispositioned. This definition is consistent with recent Cryo-ET data showing that Alphavirus budded virions can have defects at one pole of the particle^18^.

As shown in Supplementary Table 1, reducing the ED size while keeping the CTD–core fixed at 74 k_B_T resulted in a drop in the success rate to approximately 57.1%. Eliminating the TMD–TMD interactions while maintaining the WT ED size also reduced the success rate to about 28.6%, highlighting the importance of attractive interactions between spike proteins. When both the ED size was reduced and the TMD–TMD interaction was removed, the success rate declined even further to 14.3%. These results suggest that decreasing ED size disrupts assembly, likely by impairing the induction of membrane curvature, and that the absence of TMD–TMD attraction further compromises proper assembly. Our findings indicate that both steric and attractive interactions among spike proteins are essential for efficient and robust assembly. These simulations reflect the behavior of the mutant ITS described in the experimental section.

We next investigated how changes in the CTD–core interaction strength affect assembly outcomes in the mutated systems described above. Achieving the right balance is critical: an overly strong attraction can lead to excessive binding of spike proteins to the core, while a too weak interaction may fail to bring in enough spikes for successful assembly. To explore this, we performed additional simulations in which CTD–core strength was varied while holding the other parameters fixed at their mutated values. This allowed us to assess how changing the interaction strength affects budding efficiency under each mutation scenario.

Supplementary Table 2 summarizes these findings. In the absence of TMD–TMD interaction, increasing CTD–core strength from 74 to 80 k_B_T improved the success rate to 85.7% relative to WT. When only the ED size was reduced and the CTD–core was set to 75 k_B_T, the success rate reached 71.4%. However, when both the ED size was reduced and the TMD interaction was removed, success fell again to 57.1%. Notably, while stronger CTD–core attraction improves assembly efficiency, it also results in excessive spike adsorption in the WT system, underscoring the importance of a finely tuned interplay among all interactions for successful T=4 structure formation.

### Budding of an Imperfect Icosahedral Core Through the Membrane

Recent experimental studies have shown that the CHIKV capsid does not exhibit perfect icosahedral symmetry during budding^2^. To investigate this effect, we constructed a core missing one pentamer, as shown in Figure 1D, introducing an imperfect geometry. As described later, we tested this core under both wild-type conditions and in simulations incorporating mutations that disrupt CTD–core and spike–spike interactions.

Compared with the perfect icosahedral core, the imperfect core yielded a lower success rate of budding. Nevertheless, under optimized conditions—using ED = 1.15, a CTD–core interaction strength of 77 k_B_T, and a TMD-TMD = 0.1 k_B_T —spikes near the site of the missing pentamer remained correctly positioned (Figure 3), which shows the budding of an imperfect core through the plasma membrane. Reducing the ED size or weakening the TMD–TMD interactions disrupted the T = 4 symmetry by impairing inter-trimer contacts, further lowering assembly efficiency. For the optimal values of the interaction energies, only 2 out of 12 simulations produced a successful configuration (∼17%), and the success rate declined further when both the ED size and the TMD–TMD interactions were reduced.

**Figure 3.**
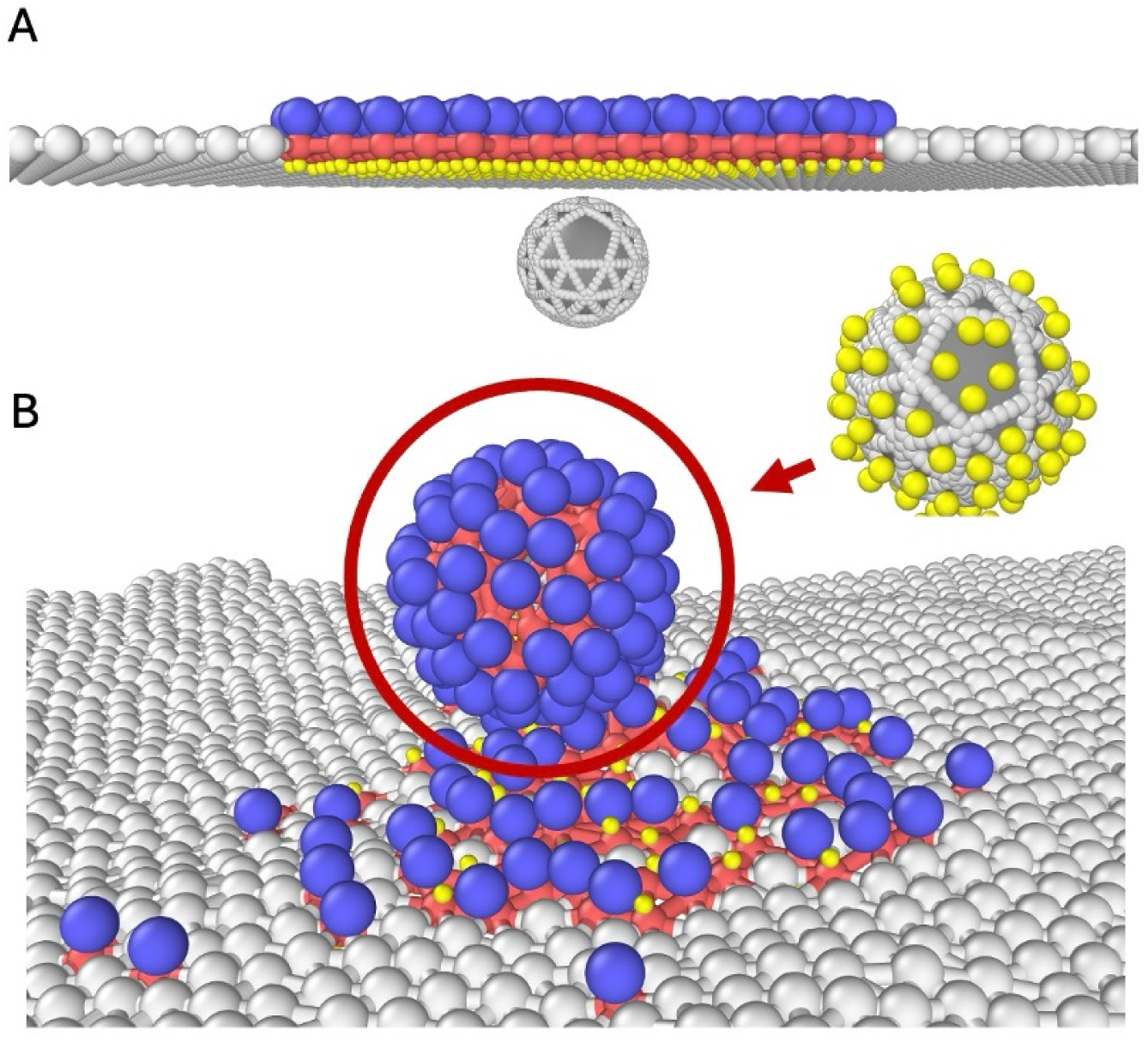
Configurations of the membrane and an imperfect core with icosahedral symmetry. (A) Initial configuration of the system with an imperfect core. (B) Final configuration after budding; the inset shows the CTD particles together with the core. System parameters: CTD–core interaction strength = 77 k_B_T, TMD–TMD interaction strength = 0.1 k_B_T. Particle diameters are 1.15 for ED, 0.8 for TMD, and 0.4 for CTD.

The relatively low success rate observed in our simulations, compared to experimental results^2^, likely stems from the limitations of the current model, which has a fixed core size and immobile subunits. Experimental data suggest that the initial core is smaller than the final assembled structure^2^, indicating that core subunits can locally dissociate and reassociate to achieve near-perfect icosahedral symmetry. In this study, our goal was to determine whether spike proteins can still assemble into an icosahedral shell in the presence of core defects, and to evaluate the roles of spike–spike and spike–core interactions in this process.

Capturing the dynamic remodeling of the core—from an imperfect to an icosahedral symmetric structure—requires a model that explicitly allows internal rearrangement of capsid subunits^44^. Simulating this process is computationally intensive, as it involves dissociation and reassociation events that must overcome significant free energy barriers^45^. However, such local restructuring is essential for correcting geometric imperfections during budding and achieving stable icosahedral symmetry. In future work, we plan to develop models that incorporate core flexibility and explicitly examine the role of the genome^46^ in promoting protein rearrangement and structural maturation.

While the theoretical model and simulations provide mechanistic insight into how specific interactions contribute to CHIKV assembly, they necessarily simplify certain aspects of the biological system, such as membrane heterogeneity and the dynamic behavior of viral proteins. To assess the biological relevance of our findings and examine how these interactions function in a cellular context, we next designed targeted mutational experiments to test the roles of glycoprotein–glycoprotein and glycoprotein-core interactions in budding and infectivity.

### The infectivity of CHIKV depends on multiple interactions between glycoproteins

To complement and validate the simulation predictions, we constructed three CHIKV mutants. To disrupt intertrimer interactions, we removed a glycosylation site in E2 via the mutation N263Q (intertrimer surface mutant, or ITS), and to weaken spike–core interactions, we introduced a double mutation in E2 Y400K/L402R (cytoplasmic tail mutant, or CT). To disrupt the intradimer (E1/E2) surface, we introduced a glycosylation site in E1 at K245N/R247T (intradimer surface mutant, or IDS). The ITS and CT mutants correspond to interaction changes examined in our coarse-grained simulations. The effect of the IDS mutation was not studied in the simulations but was included experimentally to further test the role of spike–core coupling.

Figure 4 shows a Cryo-EM reconstruction (PDB 3j2w) of the asymmetric subunit of CHIKV, depicting the positions of amino acids targeted for mutation. To investigate the role of the E1-E2 heterodimer interface, we generated an E1 double mutant K245N/R247T (CHIKV IDS) (Figure 4A) that creates a glycosylation site at the interface between E1 and E2. This is expected to introduce steric hindrance that weakens and/or destabilizes the E1/E2 heterodimer. For other Alphaviruses, it has been shown that replacing the hydrophobic amino acids in E2 that interact with the core with basic residues can inhibit budding^26, 47^. To this end, we generated the E2 double mutant Y400K/L402R to block the interaction between the core and the glycoproteins (Figure 4B) (CHIKV CT). Finally, we mutated N263 to a Q in E2 (CHIKV ITS) to remove the glycosylation site at the outer part of the trimer of heterodimers (Figure 4C). The goal was to decrease the interaction between trimers by removing the carbohydrates at this interface; we hypothesize that these carbohydrates stabilize the trimers through steric repulsion. Figure 4D is a schematic representation of the three interactions studied here, and Figure 4E depicts some of the most likely effects of the mutations on these interactions.

**Figure 4.**
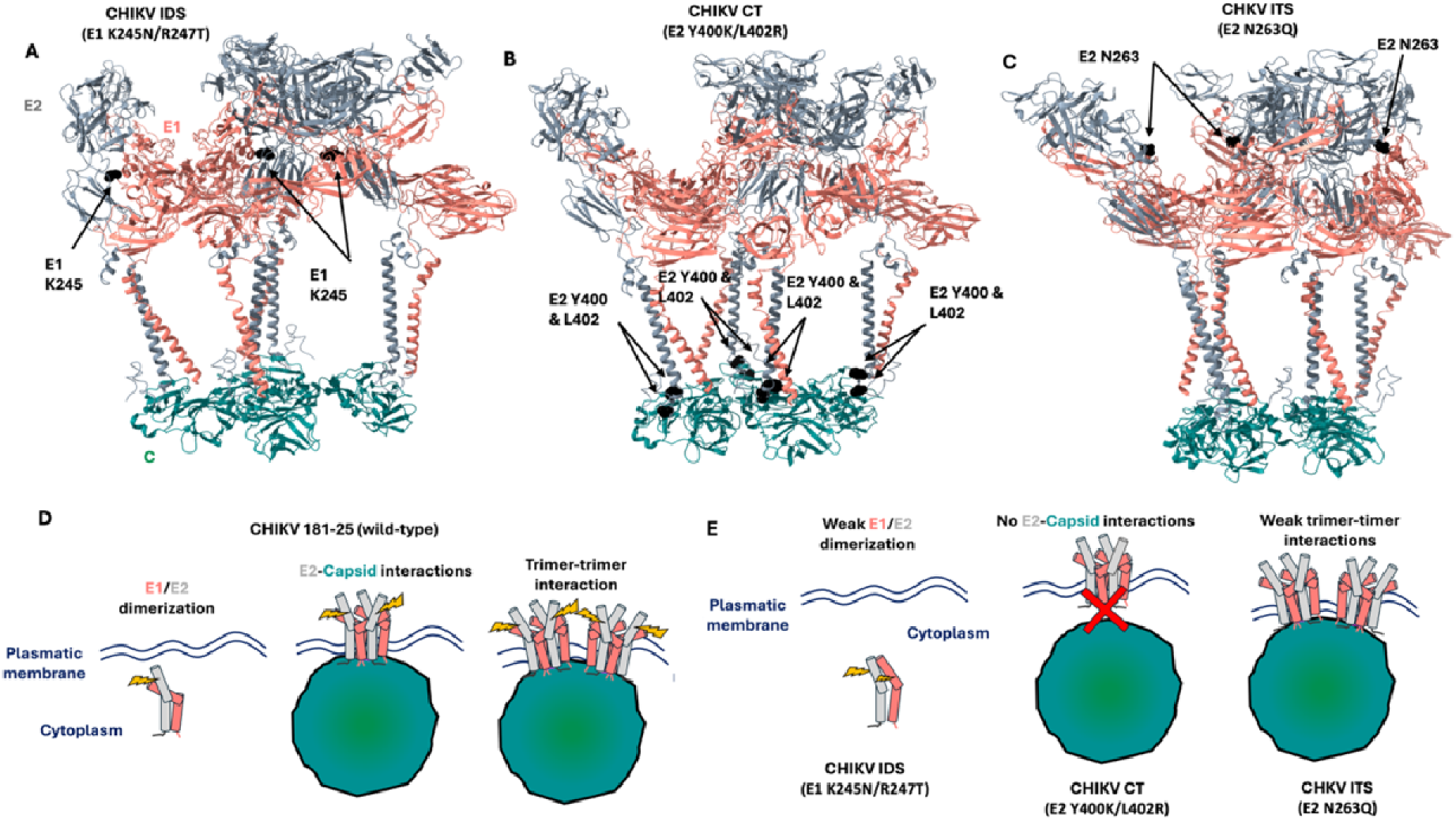
Structure of Chikungunya virus. A-C) Cryo-EM (PDB 3j2w) structure of the asymmetric unit of CHIKV virion with E1, E2, and C in salmon, gray, and teal. A) E1 K245 is at the interface between E1 and E2. B) The association of E2 with C is driven mainly by a hydrophobic pocket of E2 and Capsid. C) E2 N263 is glycosylated and might result in steric interactions between two E2 proteins at the trimer-trimer interface. D-E) Schematic representation of the interaction between glycoproteins and the core during assembly and budding. D) E1 and E2 dimerize in the cytoplasm, and once the glycoproteins are at the plasma membrane, they interact with the CHIKV cytoplasmic core, then the neighboring trimer interact with each other through high-molecular weight sugars at the E2 N263 site, E) In CHIKV IDS, a new glycosylation site was introduced in E1 amino acid 245; this mutation aims to decrease the E1/E2 heterodimer formation. CHIKV CT was designed to inhibit the interaction between E2 and the core by removing the hydrophobic pocket in E1 Y400/Y402. In CHIKV ITS, the N263 glycosylation site wa removed to weaken the trimer-trimer interactions.

### Determining the relative infectivity of mutants compared to WT using a reporter virus system

To determine the effect of each mutation on virion assembly and budding, we transfected HEK-293T cells with CMV-driven CHIKV plasmids based on the attenuated strain CHIKV 181/25 (hereafter referred to as WT). These plasmids contain a reporter gene between the C and E3 genes, which is post-translationally cleaved. Therefore, the amount of reporter gene is proportional to the number of viral structural proteins. Then, HEK-293T cells were infected with the virus rescued by transfection. At 24 hours post-infection (h.p.i.), the amount of virus produced was determined by fluorescence microscopy as a relative infectivity to the WT. Each infection was done in triplicate and with four biological replicas.

Figure 5A-D shows representative fluorescence images of infected cells (mKate2-positive). Figure 5A shows that at 24 h.p.i. most cells are mKate2-positive cells (WT infected), while panels B, C, and D show that the number of infected cells with CHIKV IDS, CT, and ITS is extremely low compared to the WT.

**Figure 5.**
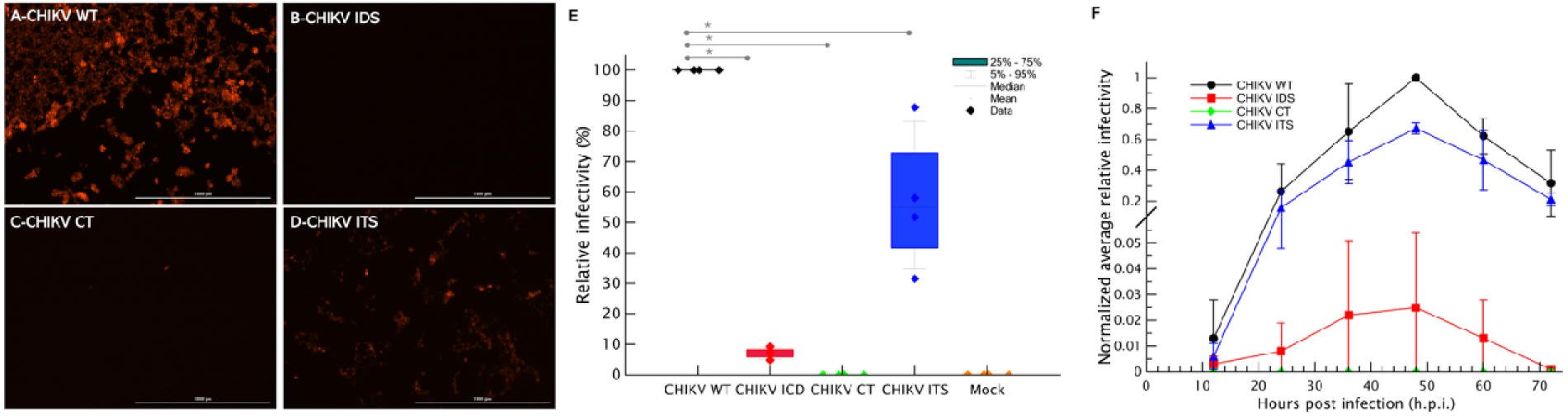
The infectivity of the viral particle depends on several protein-protein interactions. Representative fluorescent micrographs of HEK-283T cells infected with WT, IDS, CT, and ITS CHIKV viruses. The genome of all these viruses has mKate2 as a reporter gene between the capsid and E3. A-D) Cells infected with the WT, IDS, CT, and ITS viruses, respectively. E) Relative infectivity of each mutant with respect to the WT virus. The medians represent values from four experiments (n=4). K-W analysi with Dunn’s post hoc showed differences in infectivity for the three mutants compared to the WT. The mutants showed a decrease in the relative infectivity of approximately 50%, 90%, and 100%, respectively, for ISD, CT, and ITS (* p<0.05). F) Viral infection kinetics. Infections with WT, IDS, and ITS reach a maximum at 48 hours post-infection. The bars indicate the standard deviation.

The relative infectivity of these mutant CHIKV viruses is shown in Figure 5E. CHIKV IDS and CT are about 90% and 100% less infectious than the WT, respectively, while CHIVK ITS is about 50% less infectious than the WT. These results show that blocking the interaction between the core and E2 (CT) reduces relative infectivity to below the detection limit. It has been proposed that E2-core interaction is necessary for assembly ^47^. However, the data in Figure 5E suggest that this interaction is insufficient; altering, at least, the interactions between monomeric E1 and E2 and between trimers also decreases the infectivity.

These data suggest that the CHIKV mutants exhibit lower infectivity than the WT, likely due to defects in assembly and/or particle release. However, it is also possible that these mutations affect genome replication. To address this, infections with CHIKV WT, IDS, CT, and ITS were monitored over 72 hours, with measurements taken every 12 hours. Figure 5F shows the normalized average area of infected cells from three independent experiments. These results indicate that the peak of normalized relative infectivity occurs at 48 h.p.i. for CHIKV WT, IDS, and ITS. In contrast, CHIKV IDS and CT mutants show minimal, if any, replication. Therefore, the differences in infectivity between the CHIKV WT, ITS, and IDS are not due to differences in the kinetics of replication but are related to assembly/budding defects.

### Budding of CHIKV particles depends on multiple interactions

To determine which stage of the infectious cycle is affected by these mutations, we analyzed transfected cells by thin-section transmission electron microscopy (TEM) (Supplementary Figure5), focusing on how these mutations affect virion assembly and budding. CHIKV IDS and CT exhibit 90% and 100% reductions in infectivity relative to WT, respectively, making it extremely challenging to identify infected cells. Nonetheless, cells infected with Alphaviruses can often be identified by the presence of cytopathic vesicle type I (CPV-I), a characteristic marker observable by TEM^7^. Supplementary Figures 6A-D show the presence of CPV-I (arrows) in cells infected with the four types of viruses. Once these vesicles were found, we searched the area around the plasmatic membrane of that cell to characterize any possible assembly and or budding process.

Figure 6 shows characteristic budding events that result in the doubled-layer icosahedral viral particles. Figure 6A shows a CHIKV WT virion budding (red arrow) and an extracellular virion. CHIKV IDS has a relative infectivity of 90% and most budding particles are “stuck” in the initial budding stages, forming a group or collection of incomplete budding structures close to each other. (Figure 6B). The presence of partial virions suggests that this mutation did not completely block E1/E2 heterodimer formation. As expected, we could not find any budding particles for CHIKV IDS (Figure 6C). Figure 6D shows one extracellular CHIKV ITS particle and two still bound to the plasmatic membrane.

**Figure 6.**
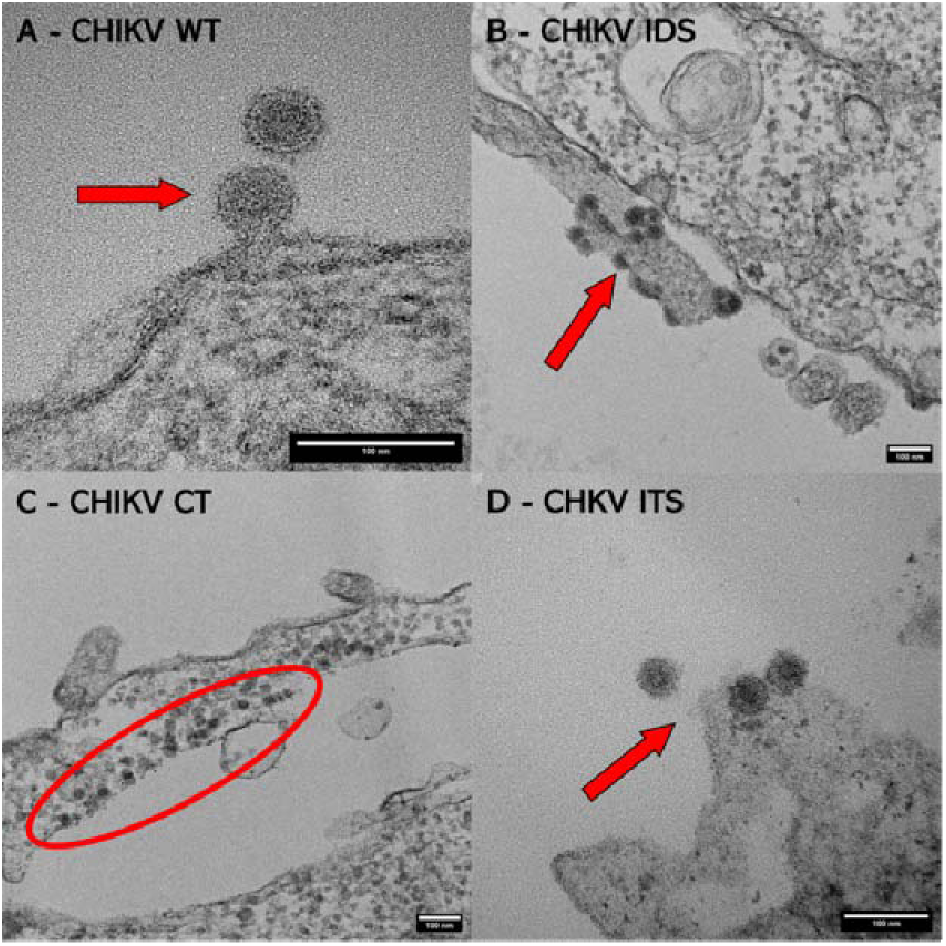
Budding structures of WT and mutant viral particles. Thin-section transmission electron micrographs of HEK-293T cells infected with the WT, IDS, CT, and ITS, CHIKV viruses showing budding events of the virions (A, B, and D). The WT, IDS, and ITS viruses bud by forming a neck, which will eventually be pinched off, releasing the virus into the extracellular medium (arrows). C) CHIKV CT does not bud out, and all the cores remain in the cytoplasm (red oval).

### Virion morphology depends on the strength of the interactions between the glycoproteins

To determine whether weakening the interaction between trimers by removing the “*molecular handshake*” between them (CHIKV ITS) alters particle morphology, we measured the hydrodynamic diameter of purified CHIKV WT and ITS particles using dynamic light scattering (DLS). Figures 7A and 7B show the hydrodynamic diameter distribution for two independent batches of CHIKV WT and ITS, respectively. The broken and solid lines indicate the measurements for batches 1 and 2, respectively. The average hydrodynamic diameters for the WT and ITS CHIKV virions are 67 nm +/- 3 nm and 113 +/- 10 nm, respectively. In addition, these experiments indicate that CHIKV ITS virions are more heterogeneous than CHIKV WT virions. This observation is consistent with the trends shown in Figure 7 and further supports our proposed mechanism. Figures 7C and D show positive-stain TEM images of purified CHIKV WT and ITS from batch 1, respectively.

**Figure 7.**
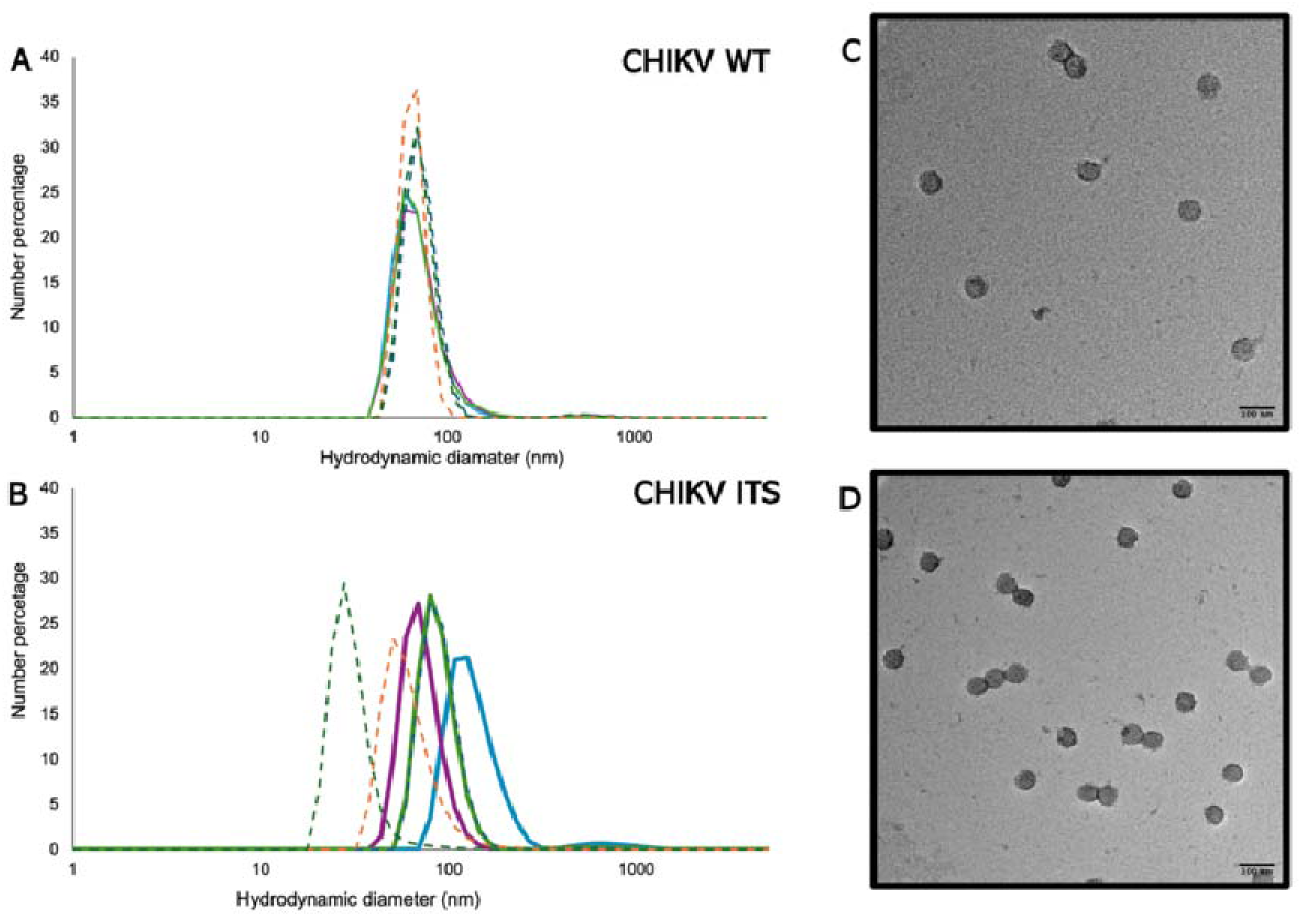
Removing the E2 N623 glycosylation site increases particle heterogeneity. The hydrodynamic diameter (nm) of purified virions was determined by using dynamic light scattering (DLS). Three measurements of the hydrodynamic diameters of the WT virus (A) and ITS (B) from two independent batches (broken lines are batch 1 and solid lines are batch 2. For the WT distribution of the hydrodynamic radius is homogenous (67 nm +/- 3 nm), while the ITS is heterogeneous (113 +/- 10 nm). C-D) Negative-staining TEM images of the purified samples for batch 1. The scale bar is 100 nm.

The data shown here suggest that altering the interaction between glycoproteins decrease the ability of the particles to bud from the producer cells. Therefore, we analyzed the cellular localization of 500 virions/cores for CHIKV WT, IDS, and ITS of transfected cells by thin-section EM. These localizations were defined as cytoplasmic, budding, and extracellular particles; the cytoplasmic particles are cores that do not interact with the plasma membrane, the budding particles are those forming any neck or protrusion, and the extracellular particles are the released virions. The distribution of virions in Supplementary Figure 7 shows that most CHIKV WT particles are either at the budding stage or released (extracellular). Very few particles remain as cytoplasmic cores. This distribution shows that when the E1/E2 heterodimer interface is altered, most particles are found at the plasma membrane at some stage of budding. This suggests that the mutant’s inability to form proper lateral interactions almost completely blocks the particle from pinching off the plasma membrane. As expected, from the WT, IDS, and ITS particles, the WT are the most abundant in the extracellular space, with only about 1/3 of the CHIKVK ITS with respect to the WT are found outside the cell, which is consistent with the relative infectivity data and the MD simulations (*i.e.,* Supplementary Table 1).

Based on our data, we hypothesize that carbohydrates at the interface between two trimers generate steric repulsion that contributes to the icosahedral ordering of the trimers. This steric interaction was first proposed for another Old-World Alphavirus, Mayaro virus, and described as a “*molecular handshake*” between the glycans of two different trimers ^31^. Therefore, removing the E2 glycosylation site at N263 is expected to decrease the stability of the viral particles. To test this, we analyzed the stability of the particles to heating by taking the supernatant from transfected cells, dividing it into aliquots, incubating it from 37 °C to 60 °C for 30 minutes, cooling it to 37 °C, and using it to infect HEK-293T cells. The relative infectivity was determined by normalizing the areas of fluorescent cells for each virus type with the sample incubated at 37 °C for that virus type (Figure 8). The normalized relative infectivity of CHIKV WT at 40°C and 45°C remained constant, whereas a significant drop in relative infectivity occurred around 50°C. CHIKV ITS is more sensitive to changes in temperature compared to a sample of CHIKV ITS incubated at 37°C. These data were fit to an equation that assumes a cooperative dissociation process. This model has two parameters: the temperature at which the relative infectivity decreases by 50% (T_RI50_) and a cooperative exponent (n). The values of T_RI50_ and n for CHIKV WT and ITS were 50°C and 99, and 44°C and 16.5, respectively. These parameters suggest that removing the glycan in E2 N263 significantly affects the particle stability and that the dissociation process of the virion of the CHIKV ITS is less ordered and might involve a weaker interaction than CHIKV WT.

**Figure 8.**
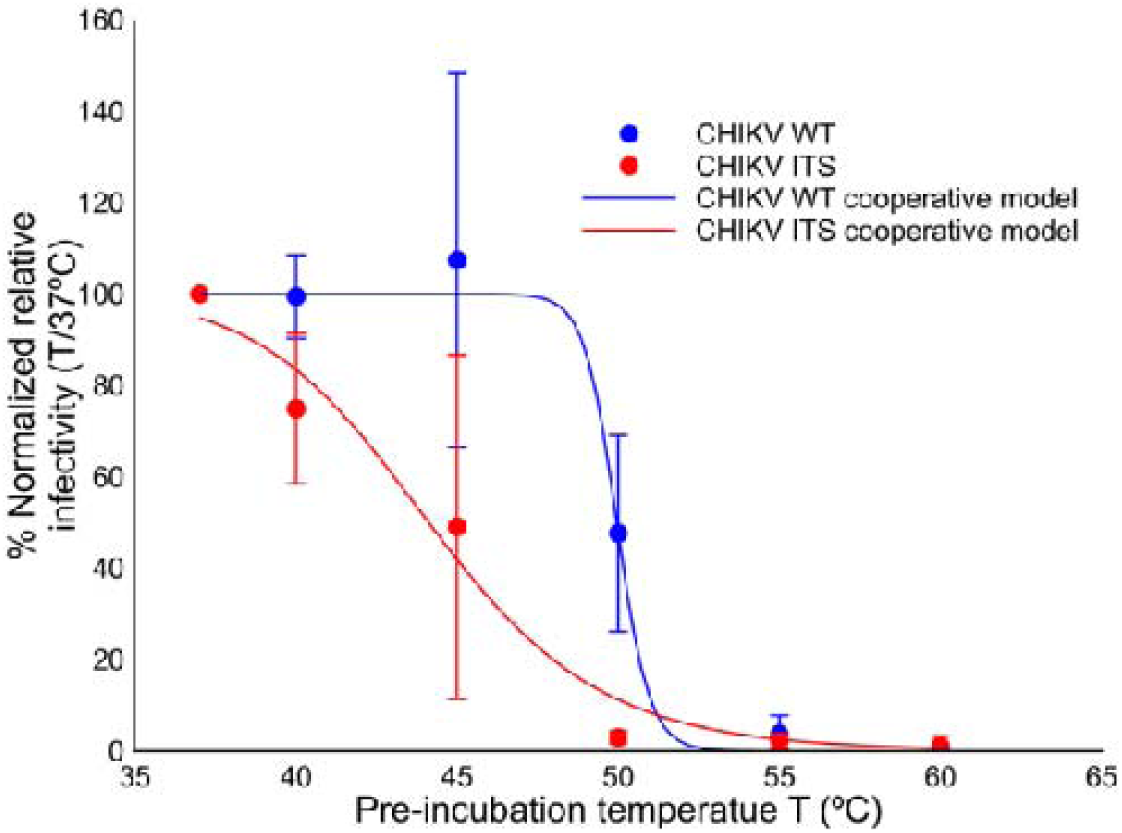
The thermal stability of the virions depends on trimer-trimer interactions. The relative infectivity of the WT and ITS viruses was calculated with respect to the area and fluorescence of infected cells for each virus at 37°C. The circles and bars are the mean and standard deviation of three independent experiments. The relative infectivity of WT remains constant from 37°C to 45°C, reaching a 50% decrease at 50°C. In CHIKV ITS decreased at 40°C, reaching a 50% decrease at 45°C. The data was fitted with a cooperative model that follows Hill’s equation (see methods and materials). For the WT, the TRI50 and n are 50°C and 99, respectively, while for the ITS mutant, they are 44°C and 16.5.

## DISCUSSION

In this paper, we set out to determine how specific protein–protein interactions govern CHIKV assembly and budding. We hypothesized that the assembly and budding depend not only on the canonical E2-core interactions ^3, 26, 35^ but also on other protein-protein interactions. To achieve this, we combined coarse-grained simulations with site-directed mutagenesis targeting key interfaces within the CHIKV glycoprotein layer.

Our simulations revealed that efficient budding requires a finely tuned balance between spike–core attraction and lateral glycoprotein interactions. Although this is not the first model to describe Alphavirus budding, it addresses novel interactions between the ectopic domains of trimers that have not been previously explored or compared with experimental data^30^.

First, interactions between the CTD of the spikes and the underlying NC were essential for initiating membrane deformation. When the strength of this interaction was reduced below a critical threshold, budding was arrested—consistent with experimental results for CHIKV CT, which disrupts the hydrophobic pocket in E2 that binds to a hydrophobic pocket on the nucleocapsid core—leading to complete loss of budding and virion assembly. The effect of this mutation has been demonstrated elsewhere^3, 26, 35^ and prompted the proposal that E2-core interactions primarily drive Alphavirus budding. However, the simulations and experiments presented here show that this interaction is necessary but not sufficient for assembly and budding.

Beyond initiating budding, interactions among glycoprotein trimers were found to be critical for achieving proper spatial organization. Simulations showed that spike–spike interactions alone could not guide self-assembly toward the correct geometry, nor could a smooth spherical core. However, an icosahedral core with T = 4 symmetry was sufficient to support the correct positioning of glycoproteins, even when imperfect. Notably, the absence of ED-ED interactions significantly reduced budding efficiency, mirroring experimental results for CHIKV ITS, in which deletion of a glycosylation site weakens trimer–trimer interactions, leading to virions with diminished infectivity, increased particle heterogeneity, and decreased particle stability. Despite a previous suggestion that spike-spike proteins between ectodomains through glycans in E2 N263 confer stability to the particle^31^, the effect of this interaction has not been previously explored either through simulation or experiments.

Although our model does not explicitly resolve E1–E2 heterodimerization, experimental analysis of the CHIKV IDS mutant—which introduces a glycosylation site in E1 at the heterodimer interface—revealed severe assembly defects. These particles were largely non-infectious, rarely completed budding, and accumulated at the plasma membrane, indicating that stable E1–E2 heterodimer formation is essential for trimer assembly and for progression beyond the early stages of envelopment and release.

High-resolution cryo-EM maps of CHIKV^2^ ^3^ □ □ □ □ □^2^ do not show electron density corresponding to high-molecular-weight glycans at E2 N263; however, our experimental data reveal a structural role for this site, consistent with the “molecular handshake” model proposed for Mayaro virus³¹. Deletion of the E2 N263 glycan disrupts trimer–trimer contacts and decreases the cooperativity of disassembly upon heating, underscoring its importance in stabilizing the spike lattice.

To further probe structural determinants of assembly, we examined how the geometry of the capsid core influences budding. Simulations performed with a defective core lacking one pentamer revealed that budding can still occur under favorable conditions, but with markedly reduced efficiency compared to a fully symmetric core. These results align with cryo-ET observations showing that one pole of the CHIKV virion can initiate budding despite local structural irregularities, emphasizing the need for geometric compatibility between the nucleocapsid and the glycoprotein lattice.

Previous work demonstrated that glycoprotein spikes alone cannot reliably form icosahedral structures³, whereas some studies suggest that capsid proteins can assemble into T = 4 shells before budding¹ ¹ . However, the symmetry of such pre-budding cores remains uncertain. Recent cryo-ET analyses¹ show that when the core deviates from perfect symmetry, spike–core and lateral spike–spike interactions can still guide assembly toward icosahedral order. These observations, together with evidence that the CHIKV core transitions from an irregular cytoplasmic structure to a more symmetric state during budding², suggest that local rearrangements of capsid subunits are required for forming stable icosahedral particles. Although our current model assumes a rigid, preassembled core and cannot capture this dynamic behavior, incorporating such flexibility in future simulations—alongside ongoing site-directed mutagenesis experiments of the core—will be crucial for elucidating the coupling between nucleocapsid remodeling and membrane budding.

## CONCLUSIONS

Our integrated computational and experimental study demonstrates that CHIKV assembly and budding are governed by a coordinated network of protein–protein interactions operating across multiple spatial and temporal scales. More significantly, this work highlights the power of simulation-guided mutagenesis as a rational strategy for dissecting and reprogramming virus assembly. By selectively altering key interaction interfaces, we gain deep molecular insights without the need for atomistic simulations or exhaustive mutational screens.

Disrupting E2–core attraction, E1–E2 heterodimerization, or trimer–trimer contacts impair assembly through distinct but complementary mechanisms—preventing budding, disrupting trimer formation, or destabilizing the glycoprotein lattice. These results counter the view that E2–core interactions alone are sufficient for particle formation; instead, they are necessary but not sufficient. Our simulations emphasize the essential roles of both vertical (core-glycoprotein) and lateral (inter-spike) interactions, including E1–E2 contacts and steric repulsions between adjacent trimers, in enabling budding and the emergence of T = 4 icosahedral symmetry.

While this study focused on perturbing E1–E2 and intertrimer interfaces, additional interactions—such as E1–E1′ and E2–E2′ contacts—are also likely to contribute to lattice stability and proper assembly. Introducing a glycosylation site into E1, as in CHIKV IDS, significantly reduced infectivity, further underscoring the delicate balance of interactions required for efficient particle formation.

A more comprehensive understanding of glycoprotein interactions will be essential for building a complete and predictive model of alphavirus assembly—one that incorporates mechanistic diversity across viral lineages and explains variations in budding efficiency and infectivity between Old- and New-World alphaviruses. Taken together, our findings support a unified model in which vertical and lateral interactions act in concert to shape virion formation, offering a strategic foundation for therapeutic interventions that target the assembly and release of infectious virus particles.

## METHODS AND MATERIALS

### Coarse-grained simulations

To simulate CHIKV budding through the plasma membrane, we developed a model incorporating a fluid membrane, spike proteins, and the viral core, as described below.

#### Fluid membrane

We model the biomembrane as a triangular lattice^48^, as illustrated in Supplementary Figure 1. The energy of this network includes contributions from both bond stretching and bending^49^. The stretching energy is represented by a harmonic potential summed over all bonds:

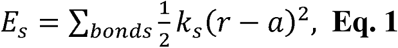

here, *a* = 10 nm is the equilibrium length of the spring. The parameter *k_s_* = 25 *k_B_T*, *nm*^-2^ is the stretching modulus. The bending energy is calculated by summing over all pairs of triangular subunits that share an edge, and is given by:

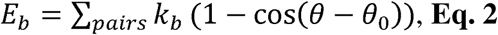

where *θ*_0_ =*π* is the preferred dihedral angle, and *k_b_* = 20 *k_B_T* is the bending modulus.

To ensure the incompressibility of our coarse-grained membrane, we have placed a spherical LJ potential particle, denoted by M (see Supplementary Figure 1), with a diameter *a* at each membrane vertex. Membrane fluidity is achieved by performing Monte Carlo (MC) simulations with the bond-flipping method, which permits dynamic changes in the membrane’s connectivity^50, 51^. The procedure for bond flips is illustrated in Supplementary Figure 2. Each MC step consists of *N_b_* =8781 attempts, where *N_b_* is the total number of bonds. In each attempt, a randomly selected bond connecting two adjacent triangles is removed and reattached between two previously unconnected vertices, as shown in Supplementary Figure 2. These moves are designed to preserve the total number of vertices and bonds, thereby maintaining the overall topology of the membrane. By allowing lateral rearrangements of connectivity, they facilitate the diffusion of topological defects across the membrane.

#### Spike proteins

Each coarse-grained spike represents a trimer of heterodimers (Figure 1). It comprises three components: the ED (ectodomain, blue), the TMD (transmembrane domain, red), and the CTD (C-terminal or cytosolic domain, yellow). ED protrudes outward from the membrane into the lumenal/extracellular-facing side of the budding virus; TMD is hydrophobic and embedded within the lipid bilayer; and CTD is located on the cytosolic side, where it interacts with the viral core.

#### Core Particles

The capsid of CHIKV is illustrated in Figure 1. In our model, we use three types of core particles. In all three types of cores, the sphere part has a diameter of 3.2a (see Eq. 1). The first is a simple spherical core (Figure 1B) that interacts with the C-terminal domain of the spike proteins (the yellow domain in Figure 1A). The second core (Figure 1C) is also spherical and interacts specifically with the CTD of the spike proteins. Its surface exhibits icosahedral symmetry and consists of 80 triangular faces. Note that only the spherical surface directly interacts with the spike proteins, while the triangles interact with them exclusively via steric repulsion. This type of interaction promotes the assembly of spikes into a T=4 shell around the core—an organization that would not occur otherwise. Without such interactions, the proteins would form a shell around a spherical core lacking any particular symmetry, which would be inconsistent with the experimental data. The third core (Figure 1D) is similar in shape and size but lacks icosahedral symmetry due to the absence of a pentamer.

#### Interactions

The attractive interactions between CTD-CTD and the core, as well as between TMD proteins, are modeled using the Lennard-Jones potential,

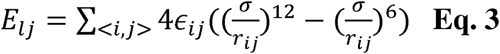

where σ= 2^-1/6*d*^ with d the distance between any two particles in contact including, CTD-core and TMD-TMD. ɛ*_ij_* and r_ij_ denote the interaction strength and the distance between particles *i*, and *j*, respectively. The interactions are subject to a cutoff distance *r_cut_t_* = 0.5*a* + *d*. In addition, the excluded volume interaction between all particles is presented with a shifted LJ repulsion potential

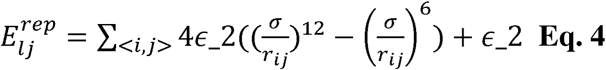

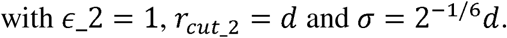

### Generation of cDNA constructs for mutant viruses

The plasmid pACNR-CHIKV is a complete sequence clone of the genome of the Chikungunya 181/25 virus (WT) in a CMV-driven plasmid^41^. It allows for the generation of viral progeny by transfecting HEK-293T cells and serves as a template for all mutations. This plasmid contains the mKate2 reporter gene between CP and E3, which is released from the structural polyprotein. The reporter gene is translated from the same subgenomic RNA as the structural proteins; hence, there is a 1:1:1:1:1 ratio of CP:mKate2:E3:E2:E1 proteins in the cytoplasm. This protein allows for the relative infectivity of the mutant viruses to be determined by fluorescence microscopy compared to the WT virus.

The intradimer surface mutant (IDS) carries two mutations in E2 (Y400K/L402R), and the intertrimer surface mutant (ITS) has N263 of E2 replaced by Q. The mutations were introduced by site-directed mutagenesis using the protocol by Liu & Naismith^52^. All PCRs were performed using HS Phusion II (ThermoFisher, Inc.) (see Supplementary Table 3). The PCR products were digested with DpnI (ThermoFisher, Inc.) and analyzed by agarose/TBE gel electrophoresis. *E. coli* DH5a bacteria (New England Biolabs) were transformed with the DpnI-digested PCR products. The selected colonies were cultured in LB (Luria-Bertani) medium with ampicillin, following standard protocols. The clones were purified with Promega’s MiniPrep Wizard kit, and the sequence identity was confirmed by Sanger sequencing at LANBAMA/IPICyT.

### Virus rescue through transfection

The production of infectious and quantification of viral particles is similar to that done with other viruses with reporter genes ^53, 54^. HEK-293T cells (ATTC) were grown in DMEM medium (Corning, Inc.) with glucose (1 g/L) and sodium pyruvate (110mg/L) with 10% fetal bovine serum (Sigma-Aldrich, Inc.) at 37°C and 5% CO_2_. The transfections were done following the protocol from Colunga-Saucedo et al.^41^. Briefly, transfections were performed in 12-well plates using Lipofectamine 3000 (Invitrogen, Inc.) according to the vendor’s protocol, with the modification that 0.6 μg of plasmid was used to transfect 100,000 seeded cells. We always transfected pACNR-CHIKV (positive control) and a mock transfection with lipofectamine, without any plasmid, on the same plate as the mutants. At 24 hours post-transfection (h.p.t.), the supernatant was removed, the cells were washed with PBS, and at 48 h.p.t., the supernatant was collected, filtered using a 0.22 μm syringe filter, and stored at -80°C in 0.1 mL aliquots. We performed four independent transfections.

### Assessing infection via a reporter assay

The quantification of the viral particles produced by transfection was directly determined as the relative infectivity of the mutants to the WT. This was achieved by infecting HEK293T cells in 24-well culture plates in triplicate with 30 μL of supernatant from transfected cells. Every culture plate used for relative quantification assays contained a positive control (CHIKV 181/25) and the supernatant of non-transfected cells (negative control), both of which were produced in the same transfection as the mutants of interest. The infectivity assays were performed only with supernatants from the same batch of transfected cells, thawed only once. The relative infectivity was measured by quantifying the total area of cells expressing mKate2 at 24 hours post-infection (h.p.i.) using an inverted fluorescence microscope LionHeart FX (Biotek, Inc.). 25 images were taken per well (5x5 symmetrical grid automatically generated by the microscope) with a 4x objective, using the Texas red filter (excitation 559 nm +/- 34 nm and emission 630 +/- 69 nm) to visualize the cells expressing the reporter gene. The relative infectivity (R.I.) was determined by measuring the area of fluorescent cells per well, averaging across wells, and setting the total average area of fluorescent cells infected with the WT as 100%. Four independent experiments (n=4) were performed, each in triplicate for each infection scenario.

### Sample preparation for thin-section EM

For thin-section Transmission Electron Microscopy (TEM), HEK293T cells were seeded in a 6-well plate, transfected, and fixed with 2% glutaraldehyde in 0.1 M cacodylate buffer, pH 7.2, for 2 hours at room temperature, following a previously published protocol^55^. The resin discs were cut (70–80 nm thickness) using a diamond blade on a LEICA EM UC7 ultramicrotome, placed on copper grids, and stained with 0.5% uranyl acetate and 0.5% lead citrate. The samples were visualized on a JEOL JEM-2100 transmission electron microscope at 200 kV with a 4k Gatan One View camera. The images were analyzed using ImageJ® (NIH) and Fiji (NIH) software.

### Viral replication kinetics

The replication kinetics were performed in three independent experiments, each in triplicate, by infecting HEK293 T cells seeded in 24-well plates (50,000 cells per well) with 15 μL of supernatant. The normalized area of the fluorescent cells was measured every 12 hours for three days, taking 25 images per well, using the Texas red filter and a 20x objective. The average fluorescent area of each experiment was normalized to WT, such that the mean fluorescence cell area of the WT was a relative infection of 100%.

### Thermal stability assays

To determine the thermal stability of the viral particles, the media collected from the transfection were diluted in OptiMem, incubated at 37°C, 40°C, 45°C, 50°C, 55°C, and 60°C for 15 minutes, and then cooled to 37°C. HEK-293T cells were infected in 48-well plates in triplicate. The R.I. of the WT at a temperature different than 37°C was normalized to that of the WT at 37°C, and the infectivity of E2 N263Q was normalized with that of this mutant at 37°C. The average R.I. was plotted as a function of the incubation temperature. The data was fitted with QtiPlot 5.15.2 using Hill’s-like equation for competitive unbinding R.I. = 100/(1+(T/T_RI50_)^n^) where R.I., T, T_RI50_, and n are the R.I. at any temperature normalized the relative infectivity at 37°C, the temperature at which the virions were incubated, the temperature at which the infectivity decreases by 50%, and a cooperative coefficient, respectively. Each experiment was done in triplicate with four biological replicates.

### Virus purification and characterization by Dynamic Light Scattering and Transmission Electron Microscopy

The production of CHIKV WT and ITS virions was performed in Vero E6 cells in two T75 flasks per virus for 3 days. The cells were grown in DMEM 10% FBS and antibiotics until they reached 100% confluency (8 million cells per flask) before infection. Then, the media was replaced with OptiMem, and the cells were infected with 1 ml of virus for each condition. The medium was replaced after 2 hours, and the media were collected, filtered through a 0.22 μm syringe filter, and stored at 4°C every 24 hours for 3 days. The viral particles were purified with a low-speed centrifugation protocol^56^. The following day, the supernatant was recovered, and the pellet was resuspended with 200 μl of PBS.

The hydrodynamic diameter of virions was measured by Dynamic Light Scattering (DLS) using a ZETA SIZE PRO Zetameter (Malvern, Inc). Each measurement was done in triplicate with two biological replicas.

The purified samples were loaded onto 100-mesh copper grids coated with Formvar (Ted Pella, Inc.) according to standard protocols, stained with uranyl acetate, and observed in a TEM (JEM-2100, JEOL) at 200 kV.

### Statistical analysis

Data analysis was performed using the Statistica 10® program. The Kolmogorov-Smirnov test was used to determine the normality of the data, and the statistical differences in the infectivity data were analyzed with the Shapiro-Wilk test. To determine whether significant differences exist (p≤0.05), the nonparametric Kruskal-Wallis test for multiple comparisons, followed by Dunn’s post hoc test, was used.

## ACKNOWLEDGMENT

M.A.C.I. (CVU: 1232434) acknowledges the support of SECIHTI through the Ph.D. fellowship 4044561 M.C-G. thanks SECIHTI for the grant CBF2023-2024-1125 and COPOCYT for the grant 2024-03-M07. RZ acknowledges support from NSF DMR-2131963 and MCB/PHY-2413062.

## ABBREVIATIONS

CHIKV: Chikungunya virus
CMV: Cytomegalovirus
CP: Capsid protein
cryo-EM: Cryo-electron microscopy
cryo-ET: Cryo-electron tomography
CT: Cytoplasmic tail mutant
CTD: C-terminal or cytosolic domain
DLS: dynamic light scattering
ED: Ectodomain
h.p.i.: Hours post-infection
IDS: Intradimer surfaced mutant
ITS: Intertrimer surface mutant
LJ: Lennard-Jones
MC: Monte Carlo
MD: Molecular dynamics
NC: Nucleocapsid or core
TMD: Transmembrane domain
TEM: Transmission Electron Microscopy
WT: Wild-type

