## Supplementary information for "Multiple protein-protein interactions drive the assembly and budding of the Chikungunya virion"

**Supplementary Material**

**Justification of** ε_mc_

When considering the interaction between the capsid and the spikes, two main components contribute: an attractive Lennard-Jones (LJ) interaction between the spikes and the spherical core, with a depth of approximately ε_mc_ = 74 k_B_T, and a repulsive excluded volume interaction between the CTD and the icosahedral core. To estimate the total effective interaction, we account for both attractive and repulsive components. Specifically, we calculate the net interaction for a single spike positioned at the center of a triangle belonging to either a pentamer or a hexamer, then take a weighted average over all such configurations to obtain the overall effective interaction energy. This yields an effective ε of approximately 24.3 k_B_T. Since each of our coarse-grained spikes represents a trimer, we divide this value by three to estimate the E2–core interaction per monomer, resulting in an effective strength of approximately 8 k_B_T, well within the range (0.9 to 9 k_B_T) used in previous models ^1^.

**Supplementary Figures**

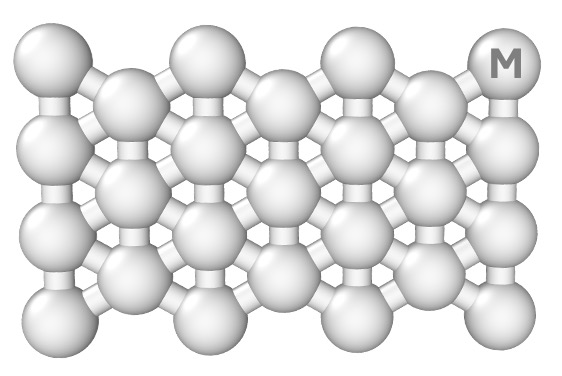

**Supplementary Figure 1.** Schematic representation of the membrane, composed of triangular elements with a bead located at each vertex. The membrane's energetics include stretching, bending, and excluded volume interactions between beads, as described by Eqs. 1–2.

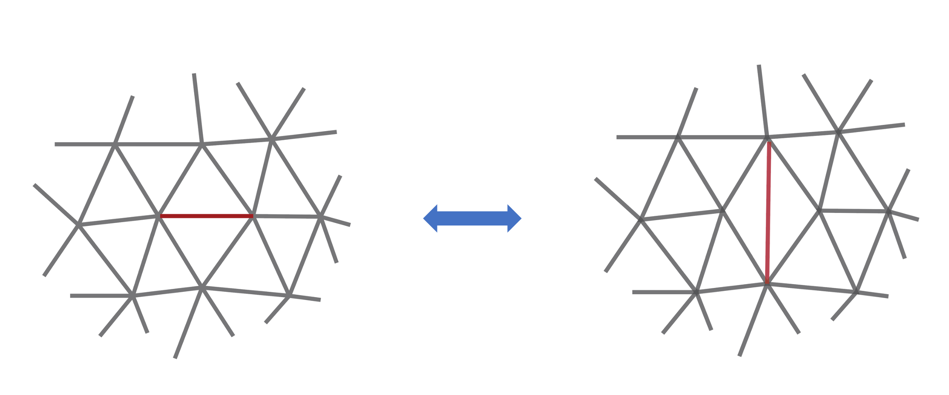

**Supplementary Figure 2**. A schematic diagram of the bond-flip method used in the MC simulations, where the shared edge between two adjacent triangles is removed and reconnected to the opposite diagonal vertices, altering the local topology of the structure.

**Budding of a Plain Spherical Particle Through the Membrane**

We observed budding even with a plain spherical core though the budding region did not exhibit perfect T = 4 icosahedral symmetry. The total number of incorporated spike proteins averaged around 79 across five simulation trials, remarkably close to the expected number of 80 for a correctly assembled T = 4 shell. This suggests that core size alone, even when commensurate with T = 4 geometry, is not sufficient to ensure correct icosahedral symmetry; in this minimal configuration, the spatial arrangement of the spikes deviates from the expected symmetry.

All relevant interaction parameters are provided in the caption of Supplementary Figure 3. These parameters were optimized to promote efficient budding, resulting in the core being decorated with ~80 spikes. We now turn to examine how the symmetry of the core influences the symmetry of the spike layer.

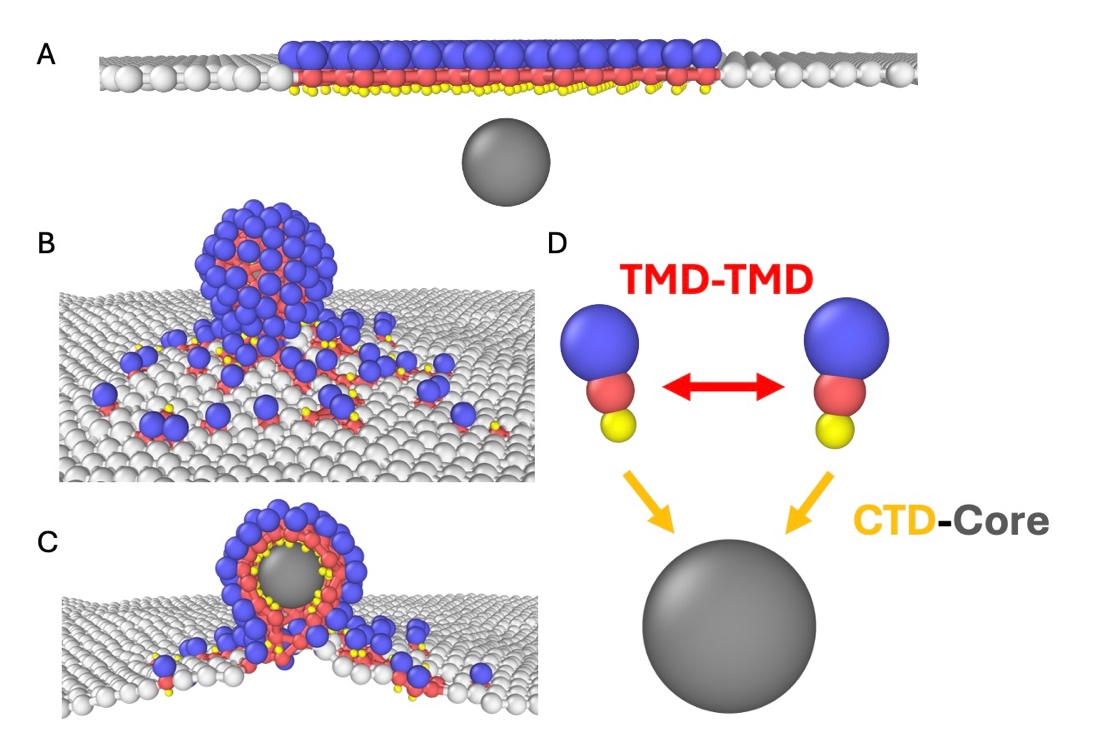

**Supplementary Figure 3**. Membrane and a plain spherical core. A) The initial state of the system before budding. B) The final state of the system after budding. The final structure contains approximately 76–82 spikes but does not exhibit icosahedral symmetry. The system parameters include a CTD–core interaction strength of 25 k_B_T and a TMD-TMD interaction strength of 0. The particle diameters are set to 1.1 for ED, 0.8 for TMD, and 0.4 for CTD. C) The cross-sectional view of the final state after budding. D) Schematic representation of TMD-TMD and CTD-core interactions. Arrows indicate attractive interactions between TMD proteins and between CTD and the core. All other particles interact through steric interactions.

**Budding of an Icosahedral Core Through the Membrane**

We now employ a core with icosahedral symmetry, as shown in Figure 1C. This core consists of 80 trimers arranged in a T = 4 structure. When the CTD–core interaction is weak, the core fails to bud from the membrane, consistent with previous studies^2-6^. We identified a critical interaction strength between the core and CTDs (ε_mc_ =15 k_B_T, see Eq. 3). When ε_mc_ falls below this threshold, the membrane remains undeformed despite the proximity of the core. This scenario captures the behavior of the mutant CT characterized in the experimental section below.

Supplementary Figure 4 A shows the initial configuration, where the core is placed near a membrane decorated with embedded spike proteins. As the simulation progresses, the attractive interaction (ε_mc_ =15 k_B_T) between the core and the CTDs drives membrane bending and initiates partial wrapping around the core. However, because the interaction strength is insufficient, budding halts prematurely, leading to incomplete envelopment (Supplementary Figure 4B). The attractive component of lateral spike–spike interactions was set to zero (see caption of Supplementary Figure 4).

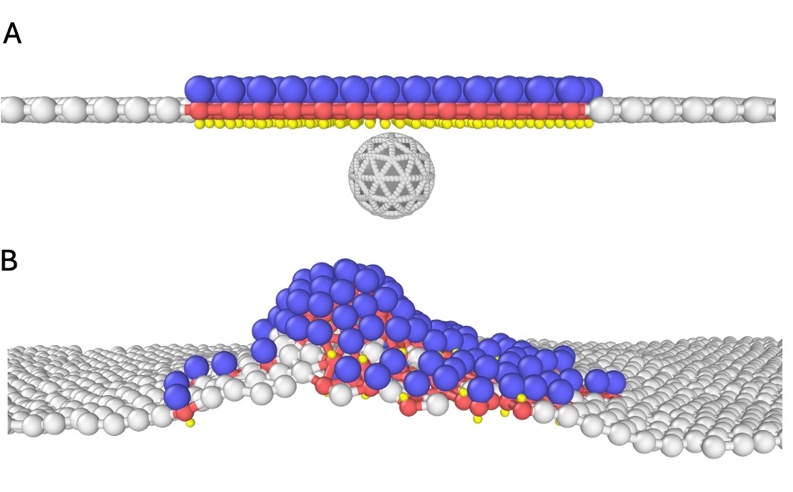

**Supplementary Figure 4**. Membrane and an icosahedral core. A) Initial configuration of the system with a core exhibiting icosahedral symmetry. B) Final configuration after introducing a CTD–core interaction with strength ε = 15 k_B_T. The core is stalled in this position. The TMD-TMD interaction strength is set to 0. The particle diameters are set to 1.1 for ED, 0.8 for TMD, and 0.4 for CTD; all diameters are in units of a (see Eq.1).

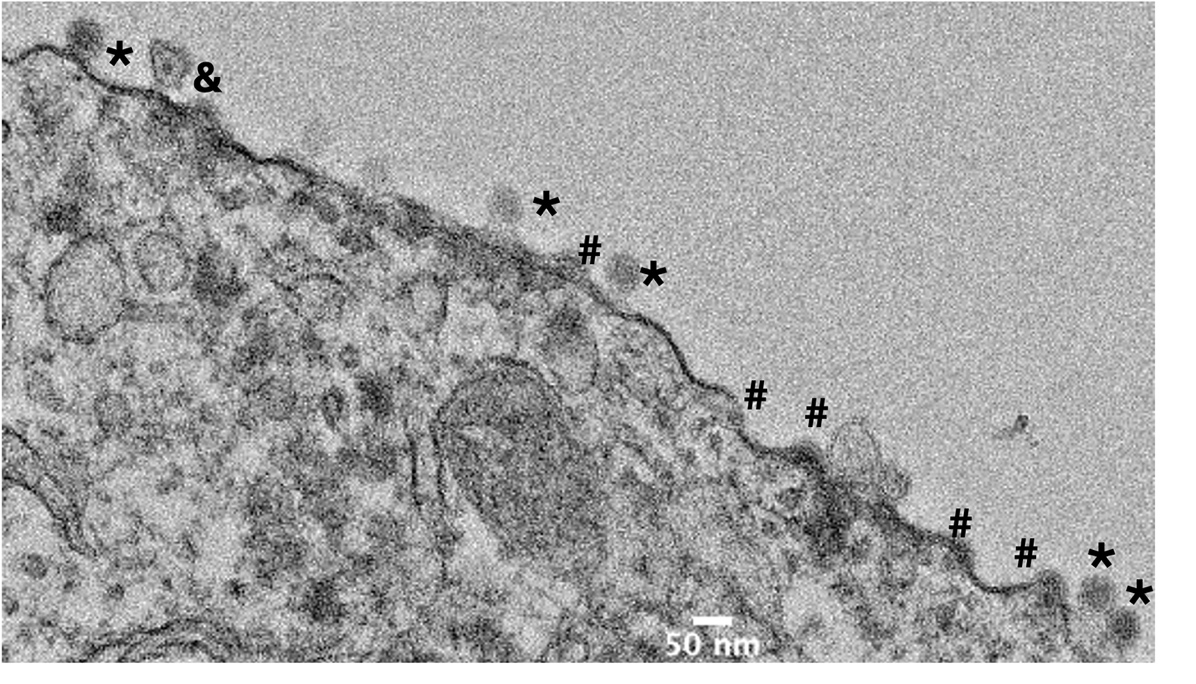

**Supplementary Figure 5.** Representative positive-stained thin-section electron micrograph of CHIKV 181-25 virions budding from HEK-293T cells. The * represents almost completely budded viral particles. The # indicates the assembly of the viral particle at the plasma membrane, and the & shows a viral RNA replication complex.

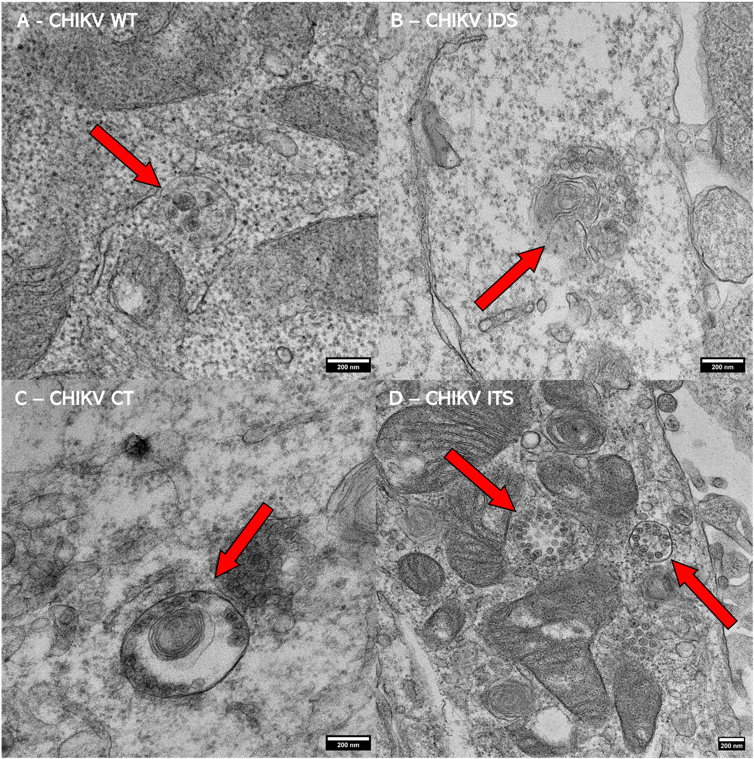

**Supplementary Figure 6.** *Cytopathic type I of infected cells.* Thin-section transmission electron micrographs of HEK-293T cells infected with the WT, IDS, CT, and ITS CHIKV viruses showing cytopathic type I (CPV-I) (see arrows) A-D) These CPV-I are characterized by small spherules surrounded by a larger membranous invagination. In the center of the spherules, a more electron-dense circular portion stands out, which is presumed to be the viral RNA. The presence of CPV-I was used to identify cells where CHIKV viral RNA is being replicated.

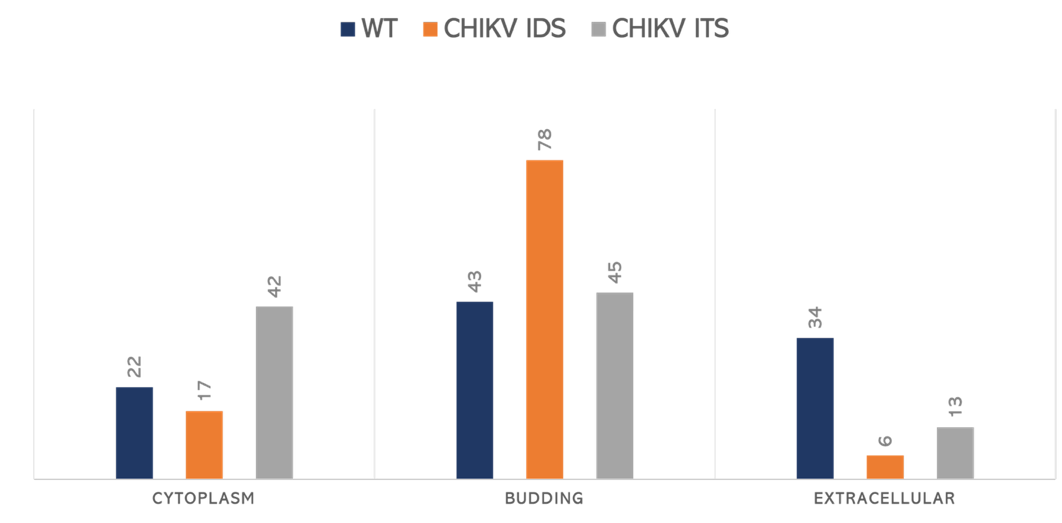

**Supplementary Figure 7.** Relative distribution (percentage) of virions in different cellular compartments. CHIKV WT (blue bar) is predominantly found in both budding at the plasma membrane and in the extracellular space. In CHIKV IDS (orange bar), which has about 2-fold fewer infections than the WT, the distribution of virions is similar to the WT. However, for CHIKV ITS (grey bar), most virions were found budding, while the amount of cytoplasmic and extracellular material was the lowest of these three viruses. These data suggest that weakening E1-E2 interactions results in defects in the budding, most likely by increasing the energy required for the core and glycoproteins to deform the plasma membrane and bud out.

**Supplementary Table 1**

| **Supplementary Table 1.** Simulation data on the Effects equivalent to mutant E2 N263. | | | |
| --- | --- | --- | --- |
| **Icosahedral core (constant CTD-Core value)** | **Parameters** | **Successful rates** | **Percentage** |
| WT | ED=1.15, CTD-Core=74, TMD-TMD=0.2 | 7/8 | 100.0% |
| Smaller ED | ED=1.1, CTD-Core=74, TMD-TMD=0.2 | 4/8 | 57.1% |
| No TMD-TMD | ED=1.15, CTD-Core=74 | 2/8 | 28.6% |
| No TMD-TMD + Smaller ED | ED=1.1, CTD-Core=74 | 1/8 | 14.3% |

For each set of parameters, we performed 8 independent simulation trials. “WT” refers to the wild-type system, which includes strong TMD-TMD interactions and an optimal ED size. The “TMD-TMD” and “CTD-Core” columns denote the interaction strengths (ε) used in the Lennard-Jones (LJ) potential. In different cases, we either reduced the ED size, removed the TMD-TMD interaction, or applied both modifications to assess their impact on the success rate. For all setups, the CTD–Core interaction strength was held constant at 74 k_B_T. The success rate is defined as the ratio of successful budding events to the total number of trials. The final column presents the success rate as a percentage relative to the WT case. “ED size” refers to the diameter of the ED particle in units of a.

**Supplementary Table 2**

| **Supplementary Table 2.** Data for the icosahedral core with varying CTD–Core interaction strength. | | | |
| --- | --- | --- | --- |
| **Icosahedral core (optimal CTD-Core value)** | **Parameters** | **Successful rates** | **Percentage** |
| WT | ED=1.15, CTD-Core=74, TMD-TMD=0.2 | 7/8 | 100.0% |
| Smaller ED | ED=1.1, CTD-core=75, TMD-TMD=0.2 | 5/8 | 71.4% |
| No TMD-TMD | ED=1.15, CTD-Core=80 | 6/8 | 85.7% |
| No TMD-TMD + Smaller ED | ED=1.1, CTD-Core=80 | 4/8 | 57.1% |

In these simulations, we varied the CTD-Core interaction and selected the value that yielded the highest success rate.

**Supplementary Table 3**

| Supplementary table 1. Primer sequences used for site-targeted mutations and sequencing. | | |
| --- | --- | --- |
| Mutant | **Primer Name** | **Sequence** |
| CHIKV ITS | FP-E2N263Q | GGCAcAaGTGACATGCAGGGTGCCTAAGGCAAGG |
|  | RP-E2N263Q | CATGTCACtTgTGCCAGAGGAAACGGAATGTGAAC |
|  | FP-SeqE2N263Q | CCAGACACCCCAGATCG |
| CHIKV IDS | FP-E1KRtNT | AAAtGAAacAGGGGCGTCGCTGCAGCACA |
|  | RP- E1KRtNT | CCCCTgtTTCaTTTAGCCAATACTTGAAGCCAG |
|  | FP-SeqE1 | TGCACCGCACACTTG |
| CHIKV CT | FP-E2Y400K | ACCGaAgGAACgGACACCAGGAGCTACCGTC |
|  | RP- E2Y400K | TGTCcGTTCcTtCGGTGTAATGCATCTGCGTCGT |
|  | RP-SeqE2Y400K | GATCACTGTTACGTGTTCG |
| Note: Lowercase letters on primers indicate nucleotides that will be changed during the site-targeted mutation. | | |
